# Dataset-Dependent Utility of Discrete Ricci Curvature for Transition-Region Ranking in Single-Cell Lineage Graphs

**DOI:** 10.64898/2026.07.25.740737

**Authors:** Chimdi Walter Ndubuisi

**Affiliations:** University of Missouri-Columbia

## Abstract

Discrete Ricci curvature is an appealing descriptor for single-cell trajectory graphs, but its practical value depends on task validity, graph-topology controls, and whether the biological target is a local transition region or a broader fate decision. We present a controlled empirical study of when curvature features add information to single-cell lineage graphs after repairing unsupported task definitions and preserving strong graph baselines. Paul15 is used as a benchmark-repair and boundary-setting dataset: on the repaired annotation-informed branch-region proxy, graph-plus-Forman and graph-plus-Ollivier improve modestly over graph topology alone (exact AUPRC 0.652 and 0.635 versus 0.607). Pancreas provides the clearest positive transition-region ranking result. On a `Fev+` endocrine transition-region benchmark, graph-plus-Ollivier reaches exact AUPRC 0.761 versus 0.669 for the graph-feature stack (five-split canonical evaluation, paired mean +0.092, 95% CI [0.076, 0.114]), and a restricted preterminal endocrine fate task improves under both curvature hybrids. The primary evaluation is transductive node ranking; graph-attachment analyses provide supporting out-of-sample robustness checks. Zebrafish provides a realism check: branch-region ranking again benefits from a hybrid model, with graph-plus-Forman strongest (0.704 versus 0.593 for graph topology), whereas a valid early `Notochord` versus `Prechordal Plate` task is graph-topology dominated on the canonical graph. Supplementary checks, including a stricter zebrafish sample-token holdout, biologically grounded bottleneck proxy, negative controls, and pairwise transfer, sharpen the same conclusion without expanding the claim set. Curvature can add useful, dataset-dependent hybrid signal for lineage transition-region tasks, but curvature-only models are weak and graph topology remains essential and sometimes sufficient.

## 1 Introduction

Single-cell lineage graphs already encode substantial information through degree, local density, centrality, connectivity, and pseudotime-like structure. Discrete curvature is therefore scientifically interesting only if it adds information beyond those graph-topology descriptors under task definitions that are biologically defensible. This paper asks a narrow question: after repairing invalid or proxy-derived tasks, when do Forman-Ricci and Ollivier-Ricci curvature features improve single-cell lineage benchmarks?

The answer is conditional rather than universal. Curvature helps most credibly on branch-region ranking tasks, where the goal is to prioritize cells in annotation-informed transition neighborhoods. The clearest Ollivier-positive result occurs on pancreas, where a `Fev+` endocrine transition region achieves the highest absolute branch-region AUPRC under graph-plus-Ollivier features. But the same evidence also rules out a universal curvature-victory narrative. Paul15 supports only a repaired, boundary-setting branch-region benchmark with modest gains; zebrafish replicates a branch-region hybrid gain while showing that a valid early-lineage task can already be solved well by graph topology.

This structure makes the study an empirical methods-and-evaluation paper rather than a geometry manifesto. The contribution is not a new trajectory engine, and it is not a claim that curvature replaces graph topology. The contribution is a controlled assessment of where curvature adds signal, where it does not, and which biological statements remain unsupported.

We make four contributions. First, we document benchmark repair on Paul15 and show why unsupported branch and fate tasks must not be used as headline evidence. Second, we compare graph topology, curvature-only features, graph-plus-curvature hybrids, random controls, rewired controls, and residualized curvature checks under a common evaluation scaffold. Third, we synthesize three datasets with different biological roles: Paul15 for boundary setting, pancreas for the clearest Ollivier-positive result and highest absolute AUPRC, and zebrafish for mixed replication. Fourth, we provide a journal-style supplementary package that makes the mathematical definitions, biological interpretation, robustness checks, and future-work boundaries explicit.

## 2 Related Work

Single-cell trajectory analysis has a mature ecosystem of graph- and pseudotime-based tools, including lineage and pseudotime methods such as diffusion pseudotime, Slingshot, PAGA, Palantir, and directed fate-mapping approaches such as CellRank (Haghverdi et al., 2016; Street et al., 2018; Wolf et al., 2019; Setty et al., 2019; Lange et al., 2022). These methods motivate the biological problem but are not the main object of replacement here. Our evaluation asks whether curvature-derived graph descriptors add information to strong graph-feature baselines on task-specific benchmarks.

The datasets are intentionally standard provenance routes rather than newly curated atlases. Paul15 is the myeloid progenitor dataset distributed through Scanpy (Paul et al., 2015; Wolf et al., 2018; Scanpy developers, 2026); pancreas is the E15.5 murine endocrinogenesis dataset distributed through CellRank and derived from single-cell profiling of pancreatic endocrinogenesis (Bastidas-Ponce et al., 2019; CellRank developers, 2026); zebrafish is the CellRank-distributed embryogenesis subset from the Farrell et al. developmental trajectory study (Farrell et al., 2018; CellRank developers, 2026). Preprocessing and graph construction use the AnnData/Scanpy data model, scikit-learn nearest-neighbor machinery, and NetworkX-style graph descriptors (Wolf et al., 2018; Pedregosa et al., 2011; Hagberg et al., 2008).

Discrete curvature has several graph formulations. We focus on the two families implemented and materialized in this repository: Forman-Ricci curvature, a combinatorial edge statistic derived from weighted cell-complex ideas (Forman, 2003), and Ollivier-Ricci curvature, a transport-based comparison of local probability measures (Ollivier, 2009). We use them as empirical feature families on rebuilt cell graphs. The paper does not assert a formal theorem connecting curvature values to biological branch points.

## 3 Methodological Framework

### 3.1 Graph construction and feature families

For each dataset, cells are nodes in a weighted nearest-neighbor graph *G* = (*V, E, w*) built from PCA coordinates. The canonical graph uses *k* = 15, Euclidean neighbor search, and heat-kernel weights

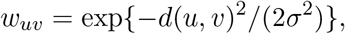

where *σ* is the median positive neighbor distance. Sensitivity analyses vary *k* and edge weighting.

The graph-feature stack includes topology and centrality descriptors: degree, weighted degree, PageRank, closeness centrality, betweenness centrality, and graph-geodesic position. The graph-geodesic position is computed as the normalized shortest-path distance from a root node selected as the minimum-weighted-degree node in the graph, using edge distances as weights. This root selection is annotation-free and does not use cell-type labels. In the out-of-sample protocol (Section 8.2), root selection and distance computation use the training graph; test cells inherit distances through their attached edges. These are deliberately strong baselines because they capture much of the available lineage signal.

### 3.2 Discrete curvature features

Forman-Ricci curvature is computed on weighted edges with unit node weights:

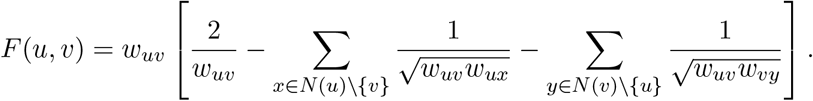

Ollivier-Ricci curvature compares lazy neighborhood measures. For idleness *α* = 0.5,

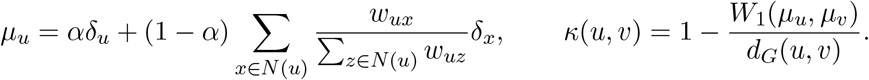

Edge curvatures are aggregated to node features using incident-edge mean, minimum, maximum, and standard deviation. We compare graph topology alone, curvature-only families, graph-plus-curvature hybrids, random matched-dimension controls, rewired graph controls, and residualized curvature controls.

### 3.3 Evaluation

The evaluation uses four tiers of increasing stringency:

#### Tier 1: Primary canonical evaluation

Logistic regression (`sklearn`, *C* = 1.0, default hyperparameters) on five stratified splits (seeds 1729–1733). Graph-derived features are computed on the full cell graph before splitting (transductive node ranking). Branch-region tasks are evaluated by exact-region AUPRC; fate-style tasks by macro-F1. This tier isolates feature informativeness from classifier capacity and provides the primary evidence for Sections 5–6.

#### Tier 2: Classifier sensitivity

Five classifiers (logistic regression, linear SVM, random forest, gradient-boosted trees, small MLP) on 20 repeated stratified splits. This tests whether curvature gains survive non-linear classifiers (Section 8.1).

#### Tier 3: Out-of-sample graph attachment

For each split, the kNN graph is built from training cells only. Test cells are attached to their *k*-nearest training neighbors, and features are recomputed on the extended graph. This reduces but does not fully eliminate test-cell influence on global graph statistics (Section 8.2).

#### Tier 4: Synthetic exact-truth benchmark

Controlled trajectories with known branch-node labels, testing curvature utility under exact ground truth (Section 8.3).

The primary inferential comparisons are Graph+Forman versus Graph and Graph+Ollivier versus Graph, reported separately. The scientifically relevant question is whether curvature adds information beyond the graph-topology stack.

## 4 Task Design, Validity Boundaries, and Supported Claims

Each dataset supports specific task definitions with explicit validity boundaries. Figure 1 and Table 1 summarize the task taxonomy.

**Table 1:**
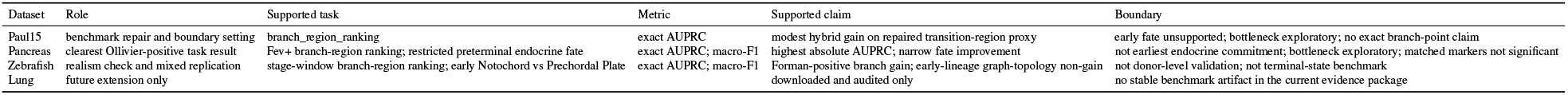
Supported task design and validity boundaries. Only biologically defensible tasks with valid label provenance are included in the primary evaluation.

**Figure 1:**
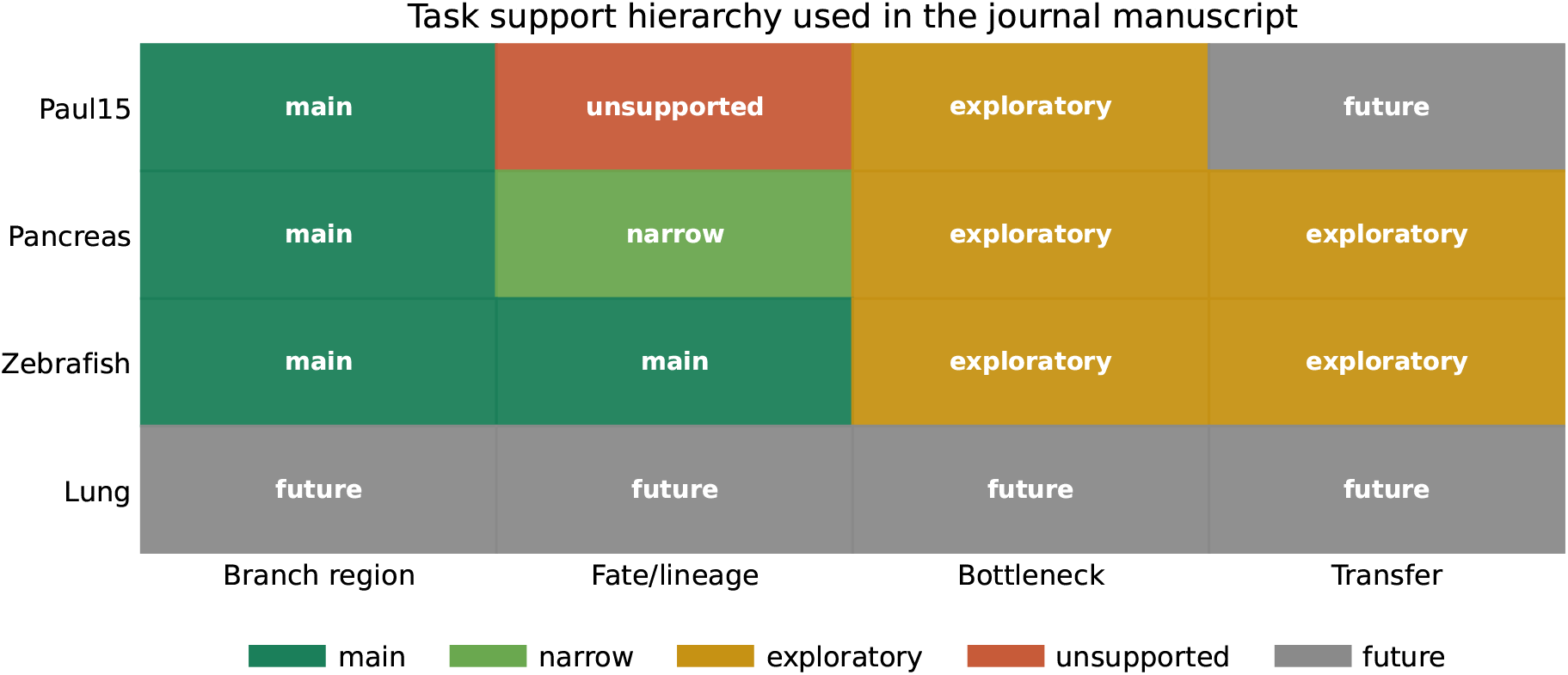
Dataset and task support hierarchy. Paul15 serves as benchmark repair, pancreas provides the clearest Ollivier-positive task result, zebrafish offers mixed replication, and lung remains future work.

Paul15 supports only repaired `branch_region_ranking` . The earlier Paul15 early-fate setup is unsupported because the split construction created unseen test classes after early-cell restriction. The repaired branch task uses annotation-informed transition-region proxy labels, not exact biological branch-point labels.

Pancreas supports two narrow but meaningful tasks: `Fev+` branch-region ranking and restricted preterminal endocrine fate prediction. The fate task is a lineage-resolution benchmark among supported preterminal endocrine states, not earliest endocrine commitment from broad `Ngn3` progenitors.

Zebrafish supports stage-window branch-region ranking and an early `Notochord` versus `Prechordal Plate` lineage task. This is the cleanest early-lineage benchmark in the current package, but it remains within-dataset rather than donor-level validation.

## 5 Results

### 5.1 Paul15 as benchmark repair and boundary setting

Paul15 is retained because it documents why task validity matters. On the repaired branch-region proxy, the graph-feature stack reaches exact AUPRC 0.607. The graph-plus-Forman and graph-plus-Ollivier hybrids reach 0.652 and 0.635, with paired exact-AUPRC deltas of +0.045 and +0.028 against the graph stack.

This supports a modest positive result: curvature is not empty on repaired Paul15. It also sets a boundary: Paul15 is not the main biological success case, does not support early fate, and does not justify an Ollivier-first or curvature-only narrative.

### 5.2 Pancreas as the clearest Ollivier-positive result

Pancreas achieves the highest absolute transition-region AUPRC. On the `Fev+` benchmark (five-split canonical evaluation), graph topology reaches exact AUPRC 0.669, graph-plus-Forman reaches 0.694 (paired Δ = +0.025, 95% CI [0.012, 0.036]), and graph-plus-Ollivier reaches 0.761 (paired Δ = +0.092, 95% percentile bootstrap CI [0.076, 0.114] over five split-level deltas, all five splits positive). The five canonical splits are not independent biological replicates; the interval reflects split-level variability rather than biological replication uncertainty.

The restricted preterminal endocrine fate task also supports a narrow fate-style gain. The graph-feature stack reaches macro-F1 0.594, while graph-plus-Forman and graph-plus-Ollivier reach 0.693 and 0.676. This does not imply broad earliest-fate prediction; it supports improved classification among preterminal endocrine states with valid class support.

### 5.3 Zebrafish as realism and non-uniform utility

Zebrafish replicates a branch-region hybrid gain but reverses the strongest curvature family. On branch-region ranking, graph topology reaches exact AUPRC 0.593, graph-plus-Forman reaches 0.704, and graph-plus-Ollivier reaches 0.641. The stronger zebrafish branch result is therefore Forman-based.

The early-lineage benchmark is the realism check. On the canonical graph, graph topology reaches macro-F1 0.771, graph-plus-Forman reaches 0.764, and graph-plus-Ollivier reaches 0.742. The paired deltas do not support a robust curvature improvement. This is a valid non-gain: graph topology is already sufficient for this broad early-lineage separation.

### 5.4 Cross-dataset synthesis

Table 2 summarizes the main supported tasks, and Figure 2 shows branch-region performance across datasets. Across datasets, three conclusions are stable. First, curvature helps most credibly on branch-region tasks. Second, the strongest curvature family depends on the dataset: Ollivier has the clearest flagship result on pancreas, while Forman has the stronger consistency story on Paul15 and zebrafish branch ranking.

**Table 2:** Compact main results on the canonical graph. Branch tasks use exact-region AUPRC; fate-style tasks use macro-F1.

| Dataset | Task | Metric | Graph | Graph+Forman | Graph+Ollivier | Best family | Delta |
| --- | --- | --- | --- | --- | --- | --- | --- |
| paul15 | branch_region_ranking | auprc_exact | 0.607 | 0.652 | 0.635 | graph_plus_forman | 0.045 |
| pancreas | branch_region_ranking | auprc_exact | 0.669 | 0.694 | 0.761 | graph_plus_ollivier | 0.092 |
| pancreas | preterminal_endocrine_fate_prediction | macro_f1 | 0.594 | 0.693 | 0.676 | graph_plus_forman | 0.099 |
| zebrafish | branch_region_ranking | auprc_exact | 0.593 | 0.704 | 0.641 | graph_plus_forman | 0.111 |
| zebrafish | early_lineage_prediction | macro_f1 | 0.771 | 0.764 | 0.742 | graph_feature_stack | 0.000 |

**Figure 2:**
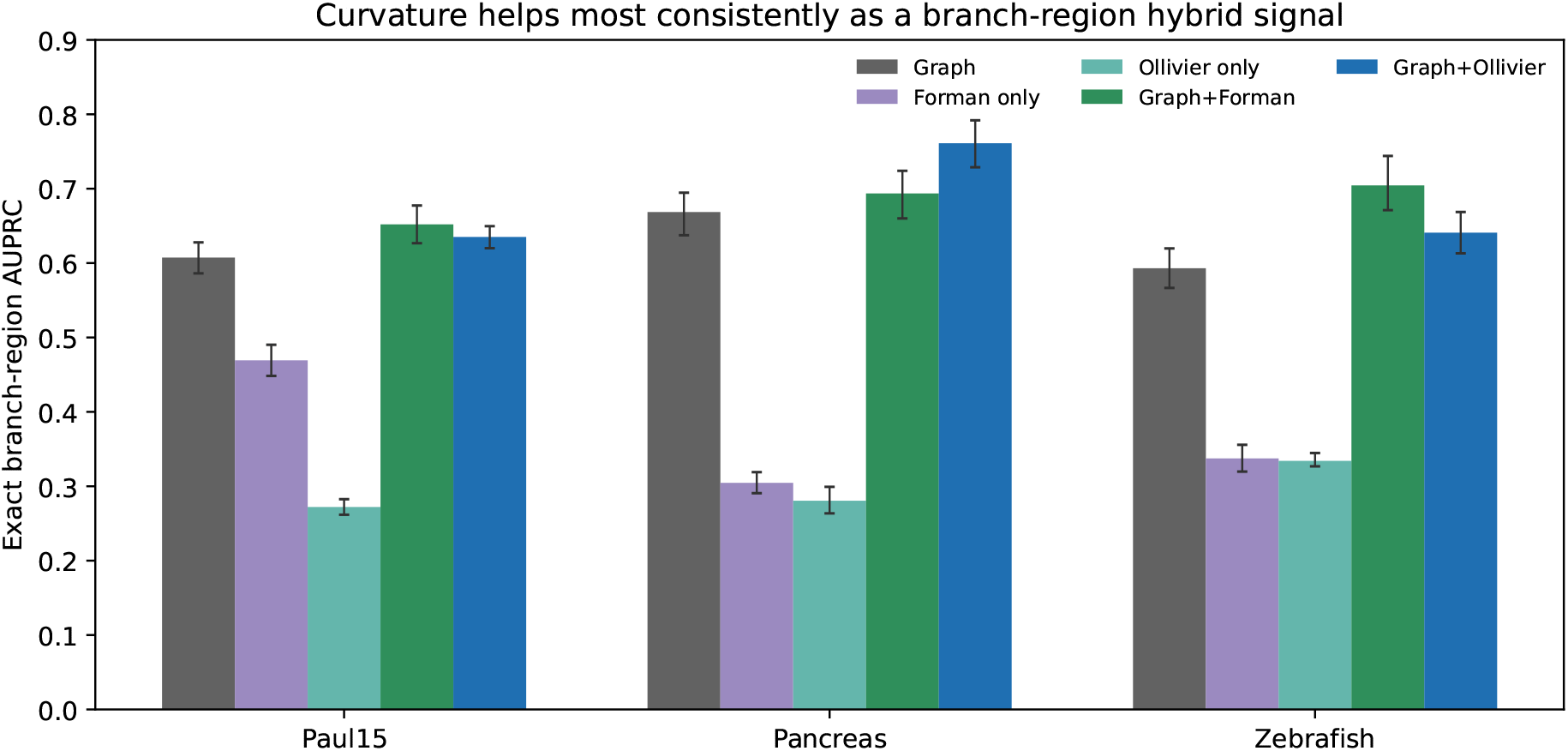
Branch-region ranking across datasets. Curvature-only models remain weak, while graph-plus-curvature hybrids improve over graph topology on the supported branch-region tasks. The strongest family is dataset-dependent: pancreas favors Ollivier, while Paul15 and zebrafish favor Forman.

Third, fate-style tasks are heterogeneous: pancreas preterminal fate benefits from curvature hybrids, whereas zebrafish early-lineage prediction does not show a canonical gain.

## 6 Biological Interpretation

The task-level interpretation is strongest in pancreas. The `Fev+` population is an annotation-backed endocrine transition region with high graph-neighborhood mixing. The graph-plus-Ollivier hybrid concentrates top-ranked predictions more tightly on exact `Fev+` cells than the graph stack: among the top 50 branch predictions, graph topology recovers 37 exact positives, graph-plus-Forman recovers 39, and graph-plus-Ollivier recovers 43.

### Marker validation

We compare endocrine transition markers between the top-50 and bottom-50 curvature-ranked cells. In the unmatched comparison, five transition markers are significantly upregulated in top-ranked cells (Chga, Pax6, Mafb, Pax4, Nkx6-1; all *p <* 0.01). However, when top-ranked cells are matched to controls on pseudotime, degree, and library size, no marker survives Benjamini-Hochberg correction (0/14 significant). This indicates that the unmatched enrichment was confounded by pseudotime and graph-position differences between high- and low-curvature cells. The curvature ranking correlates with transition biology, but the marker signal is not independent of the graph-topological features that also predict branch-region labels.

### Trajectory baseline comparison (20-repeat evaluation)

DPT alone reaches AUPRC 0.181 on pancreas, substantially below the graph-feature stack (0.648 in the 20-repeat evaluation; 0.669 in the five-split canonical evaluation). Graph+Ollivier reaches 0.749 in the 20-repeat evaluation versus DPT+Ollivier 0.269. The graph-feature stack outperforms DPT on these transition-region tasks.

Zebrafish separates local transition-region ranking from broad early-lineage prediction. The branch-window task benefits from a hybrid model, but the early `Notochord` versus `Prechordal Plate` task is already well served by graph topology. This non-uniformity is biologically plausible: datasets differ in organism, developmental process, annotation granularity, graph construction, and how much lineage structure is already captured by topology.

## 7 Robustness, Negative Controls, and Appendix-Guided Checks

Supplementary analyses refine the scope of the primary findings without expanding the claim set. Negative controls (random features, rewired graphs) remain below graph topology on branch-region tasks. Sensitivity checks show that graph construction parameters can affect magnitudes while preserving the broad branch-region pattern. Residualized curvature checks reduce concern that the pancreas branch result is only a hubness artifact, but they do not establish a mechanistic explanation.

The stricter zebrafish single-stage sample-token holdout still favors graph topology (macro-F1 0.711) over graph-plus-Forman and graph-plus-Ollivier (0.670 and 0.674). The pancreas late-`Ngn3 high EP` bottleneck proxy is biologically grounded but weak, with exact AUPRC 0.109 for graph-plus-Ollivier versus 0.096 for graph topology. Pairwise transfer is asymmetric, with moderate Paul15-to-pancreas transfer and weak transfers involving zebrafish. These supporting analyses are reported in the supplement.

**Figure 3:**
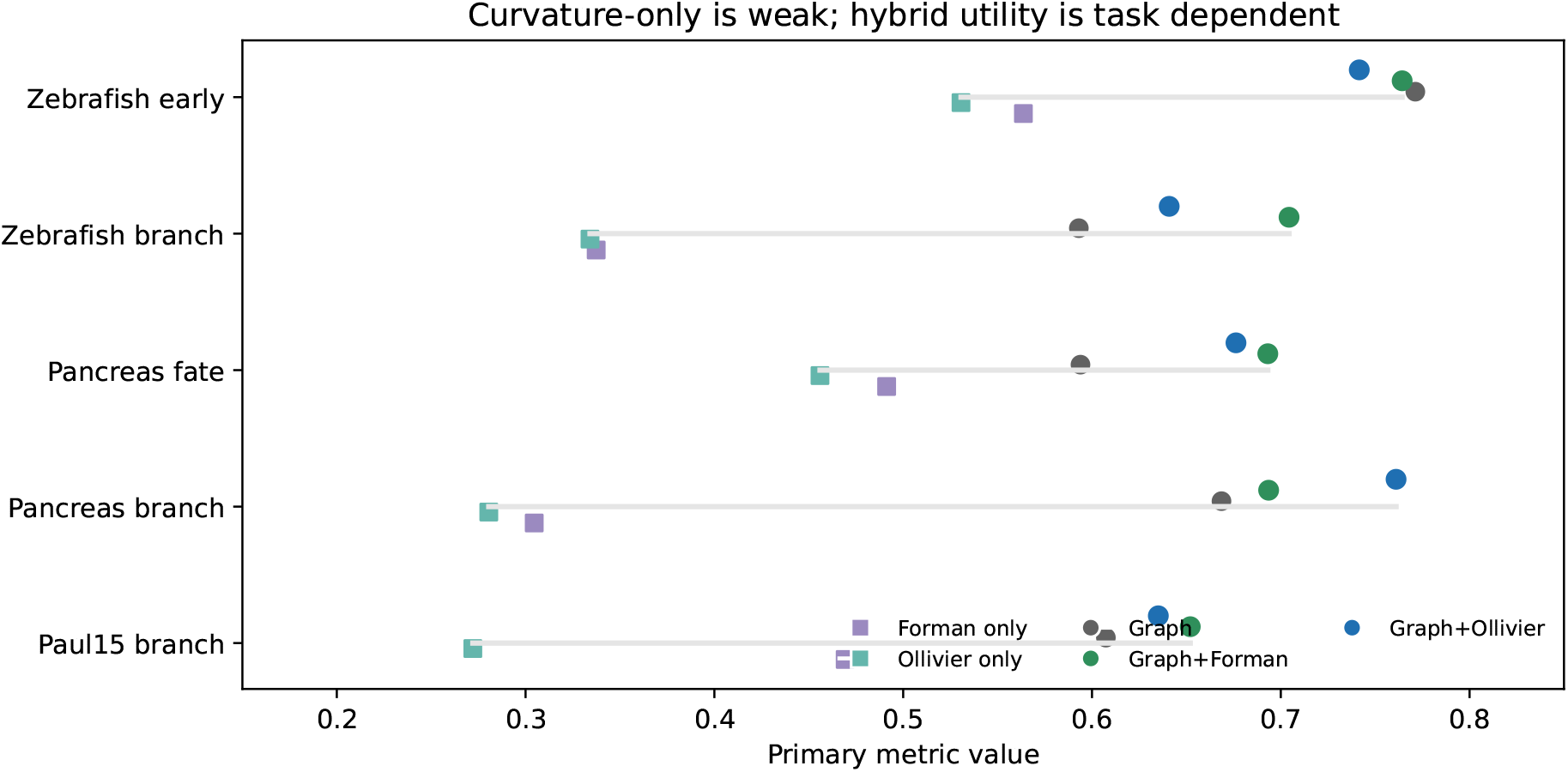
Topology, curvature-only, and hybrid feature families across supported tasks. The central pattern is hybrid and non-uniform: curvature adds signal in several transition-region settings, but graph topology remains strong and curvature-only models are not sufficient.

## 8 Sensitivity Analysis

### 8.1 Classifier sensitivity

We evaluate five classifiers on all three branch-region tasks using 20 repeated stratified splits. Table 3 reports Graph+Forman and Graph+Ollivier deltas separately.

**Table 3:**
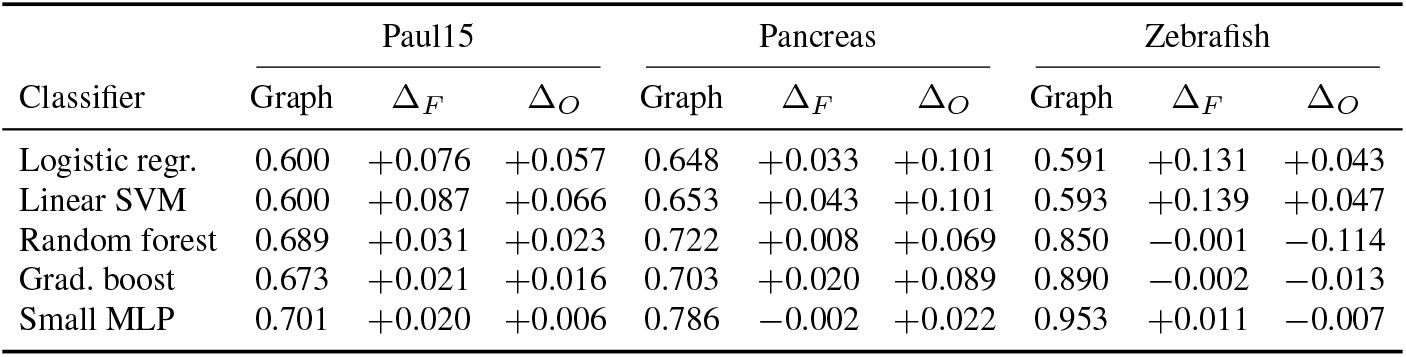
Curvature deltas by classifier and dataset (20 repeated stratified splits). Δ_*F*_ : Graph+Forman minus Graph. Δ_*O*_: Graph+Ollivier minus Graph. Prevalence: Paul15 16.6%, Pancreas 23.2%, Zebrafish 19.4%.

| Classifier | Paul15 |  |  | Pancreas |  |  | Zebrafish |  |  |
| --- | --- | --- | --- | --- | --- | --- | --- | --- | --- |
| | Graph | $\Delta_F$ | $\Delta_O$ | Graph | $\Delta_F$ | $\Delta_O$ | Graph | $\Delta_F$ | $\Delta_O$ |
| Logistic regr. | 0.600 | +0.076 | +0.057 | 0.648 | +0.033 | +0.101 | 0.591 | +0.131 | +0.043 |
| Linear SVM | 0.600 | +0.087 | +0.066 | 0.653 | +0.043 | +0.101 | 0.593 | +0.139 | +0.047 |
| Random forest | 0.689 | +0.031 | +0.023 | 0.722 | +0.008 | +0.069 | 0.850 | -0.001 | -0.114 |
| Grad. boost | 0.673 | +0.021 | +0.016 | 0.703 | +0.020 | +0.089 | 0.890 | -0.002 | -0.013 |
| Small MLP | 0.701 | +0.020 | +0.006 | 0.786 | -0.002 | +0.022 | 0.953 | +0.011 | -0.007 |

Two findings emerge. First, curvature gains diminish with more powerful classifiers on Paul15 and zebrafish, suggesting that non-linear models can learn equivalent feature interactions from graph topology alone. Second, the pancreas Ollivier signal persists under random forest (Δ_*O*_ = +0.069) and gradient-boosted trees (Δ_*O*_ = +0.089), indicating that the Fev+ transition-region signal is not merely a linear-classifier artifact. Zebrafish Ollivier gains reverse under tree-based methods (Δ_*O*_ = *−* 0.114 with random forest). The dataset-dependent pattern holds across classifiers: pancreas favors Ollivier, Paul15 and zebrafish favor Forman.

### 8.2 Shared-graph out-of-sample attachment (Protocol A)

To test whether the curvature gain depends on the transductive graph, we evaluate under out-of-sample graph attachment: for each of 10 CV splits, the kNN graph is built from training cells only, test cells are attached via *k*-nearest training neighbors, and all features are recomputed on the extended graph. This protocol ensures test cells do not participate in the original graph construction. However, after attachment, test cells can influence training-node centralities and each other’s curvature through the shared extended graph, so the protocol is not strictly inductive. Table 4 reports both Forman and Ollivier results with Holm-corrected paired Wilcoxon *p*-values.

**Table 4:** Protocol A (shared-graph attachment): training graph built from training cells, test cells attached via kNN; features recomputed on extended graph. AUPRC mean over 10 splits. Holm-corrected Wilcoxon *p*-values. Prevalence: Paul15 16.6%, Pancreas 23.2%, Zebrafish 19.4%.

|  | Paul15 |  | Pancreas |  | Zebrafish |  |
| --- | --- | --- | --- | --- | --- | --- |
| | AUPRC | $p_{\text{Holm}}$ | AUPRC | $p_{\text{Holm}}$ | AUPRC | $p_{\text{Holm}}$ |
| *Logistic regression* |  |  |  |  |  |  |
| Graph only | 0.512 | — | 0.607 | — | 0.543 | — |
| Graph + Forman | 0.658 ($\Delta+0.146$) | 0.023 | 0.645 ($\Delta+0.038$) | 0.023 | 0.623 ($\Delta+0.080$) | 0.109 |
| Graph + Ollivier | 0.584 ($\Delta+0.072$) | 0.023 | 0.657 ($\Delta+0.050$) | 0.023 | 0.566 ($\Delta+0.023$) | 0.023 |
| *Random forest* |  |  |  |  |  |  |
| Graph only | 0.593 | — | 0.545 | — | 0.643 | — |
| Graph + Forman | 0.607 ($\Delta+0.014$) | 0.750 | 0.561 ($\Delta+0.016$) | 0.393 | 0.745 ($\Delta+0.101$) | 0.023 |
| Graph + Ollivier | 0.637 ($\Delta+0.044$) | 0.023 | 0.621 ($\Delta+0.076$) | 0.049 | 0.653 ($\Delta+0.010$) | 0.750 |

**Table 5:** Cross-dataset synthesis of graph topology, curvature-only behavior, and hybrid utility.

| Dataset/task | Topology | Curvature pattern | Claim level |
| --- | --- | --- | --- |
| Paul15 branch | strong | modest hybrid gain, Forman hybrid best | main supported but boundary-setting |
| Pancreas branch | competitive | clearest Ollivier-positive hybrid gain | main supported, highest absolute AUPRC |
| Pancreas preterminal fate | usable | both hybrids improve, Forman slightly ahead on canonical graph | main supported, narrow fate claim |
| Zebrafish branch | competitive | clear Forman-positive hybrid gain | main supported mixed replication |
| Zebrafish early lineage | already strong | no clear canonical improvement | main supported non-gain |

**Figure 4:**
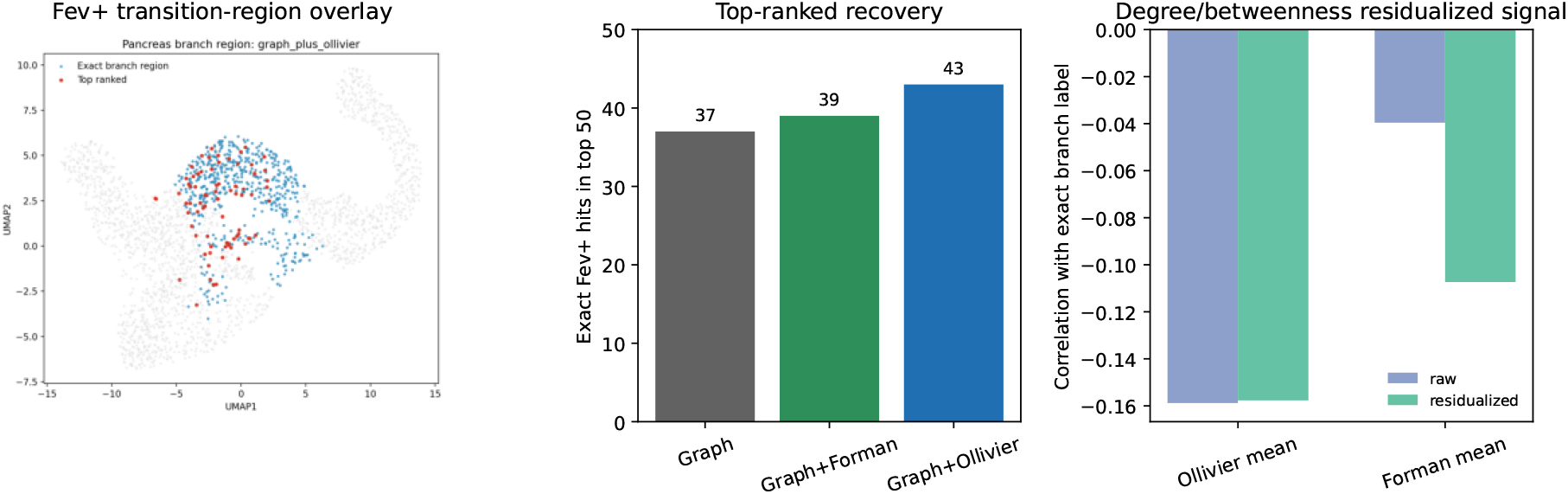
Pancreas interpretation. The clearest Ollivier-positive task result is on the `Fev+` transition-region benchmark, not exact branch-point localization. Top-ranked recovery quantifies Fev+ label retrieval and should not be interpreted as independent marker-level biological validation. The residualized panel reports curvature-signal correlations after degree/betweenness adjustment and is used only as a bounded support check.

**Figure 5:**
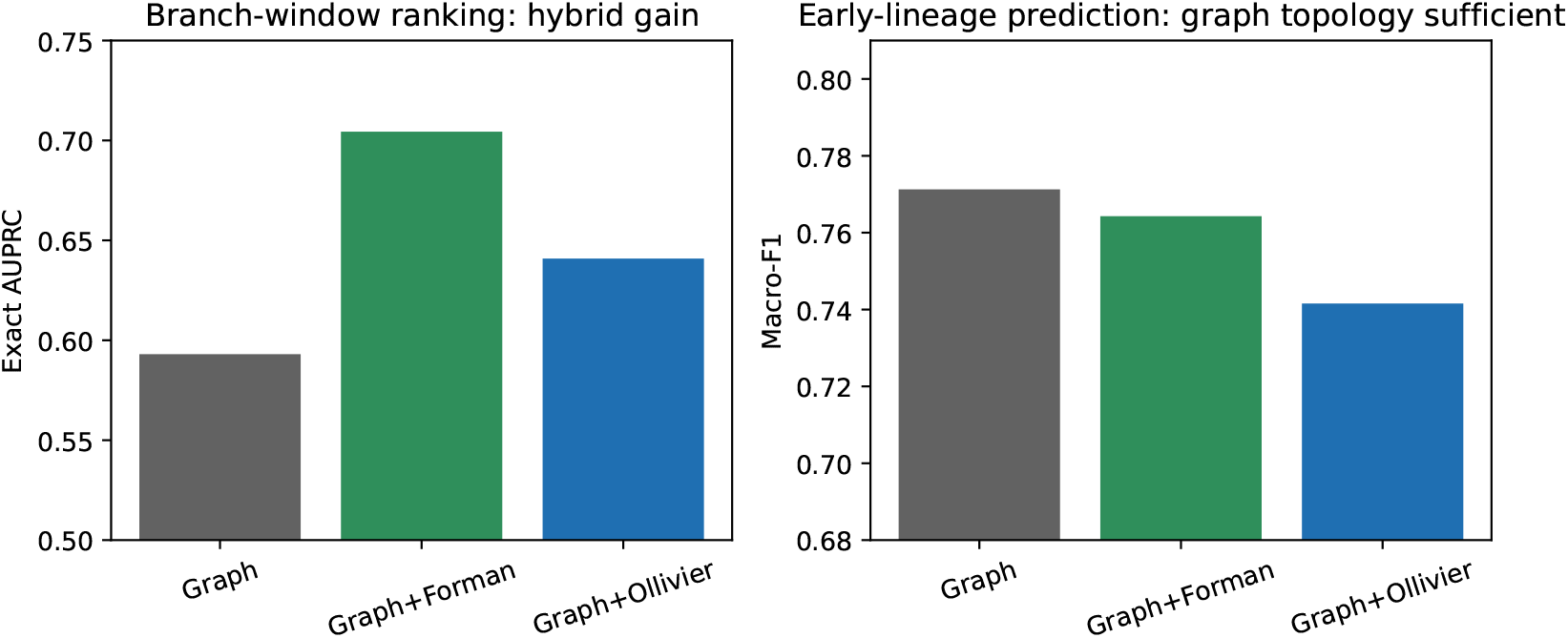
Zebrafish realism check. The same dataset supports a branch-region hybrid gain and a canonical early-lineage non-gain, showing that curvature utility depends on task geometry rather than on dataset inclusion alone.

Both Forman and Ollivier curvature gains persist under out-of-sample graph attachment with Holm-corrected statistical support. The flagship pancreas Ollivier result survives: Δ_*O*_ = +0.050 (LR, *p*_Holm_ = 0.023) and Δ_*O*_ = +0.076 (RF, *p*_Holm_ = 0.049). This reduces concern that the observed gain is solely caused by computing features on the full graph, though absolute performance is lower and the protocol does not fully eliminate test-cell mutual influence.

#### Strict frozen-training-graph attachment (Protocol B)

A stricter Forman-only analysis freezes training-node features and prohibits test-to-test edges. Forman gains persist for Paul15 (RF Δ_*F*_ = +0.091) and zebrafish (RF Δ_*F*_ = +0.235) but are mixed on pancreas (RF Δ_*F*_ = +0.057, LR Δ_*F*_ = *−*0.051). Ollivier was not evaluated under Protocol B due to computational cost, so these results do not establish strict-inductive persistence of the canonical pancreas Ollivier gain. Variance is higher under Protocol B, reflecting the harder evaluation setting. A complete strict-inductive comparison of both Forman and Ollivier remains necessary.

### 8.3 Synthetic exact-truth benchmark

We generate three synthetic trajectory scenarios with exact branch-node labels to isolate the mechanism by which curvature adds information. Each scenario embeds 2D trajectories with Gaussian noise (*σ* = 0.03), builds a kNN graph (*k* = 15, heat-kernel weights), and evaluates with 20 repeated stratified splits (25% test).

**Y-branch** (*n* = 600, 33% branch prevalence): a trunk splits into two symmetric branches. Branch nodes are cells within *±*0.08 of the bifurcation point. Graph topology alone reaches AUPRC 0.982 ± 0.012 (LR) and 0.970 *±*0.023 (RF). Neither Forman (Δ_*F*_ = 0.000) nor Ollivier (Δ_*O*_ = *−*0.008) adds signal. On balanced trajectories, graph centrality features are sufficient.

**4-way tree** (*n* = 800, 20% branch prevalence): a trunk splits into four branches. Forman adds nothing (Δ_*F*_ = 0.000); Ollivier provides a small gain (Δ_*O*_ = +0.011 LR, +0.003 RF), within noise.

**Density-imbalanced Y-branch** (*n* = 600, one branch 3*×* denser than the other, 17% branch prevalence): Ollivier curvature provides a substantial gain with random forest (Δ_*O*_ = +0.093, from 0.826 to 0.920), while Forman does not (Δ_*F*_ = *−*0.002). In this synthetic configuration, Ollivier curvature improves ranking under branch-density imbalance. An expanded sweep over 20 generative seeds, five density ratios (1.0–5.0), four noise levels, and four branch angles shows that Forman provides small positive gains (+0.02 to +0.05) across the evaluated synthetic conditions, while a nonbranching negative control shows near-zero delta (+0.0002). Broader variation is required before treating density imbalance as an established mechanism.

## 9 Limitations and Future Directions

The study has clear limits. Classifier sensitivity analysis (Section 8.1) shows that curvature gains diminish with non-linear classifiers on two of three datasets, framing curvature as a compact engineered feature most useful for linear models. The pancreas Ollivier result is the exception, persisting under all tested classifiers. Out-of-sample graph attachment (Section 8.2) reduces concern that the transductive protocol drives the gain, but absolute AUPRC values are lower and the protocol does not fully prevent test-cell mutual influence.

The primary evaluation uses transductive node ranking. Branch-region labels are proxy transition regions rather than exact biological branch-point labels. Unmatched marker validation shows significant enrichment of transition markers in top-ranked curvature cells (Section 6). However, after matching on pseudotime, degree, and library size, no marker survives multiple-testing correction (0/14 BH-significant).

The unmatched marker signal was confounded by the same graph-topological features that drive the branch-region classification.

The supported fate tasks are narrow: pancreas is preterminal endocrine fate, and zebrafish is an early two-lineage window. All evaluations are within-dataset and weaker than donor-level or external validation. Curvature-only models are not sufficient. A comparison with diffusion pseudotime (DPT) shows that DPT alone is weaker than the graph-feature stack on all three datasets (AUPRC 0.181–0.636 versus 0.591–0.648), and curvature-augmented DPT does not match the full graph+curvature hybrid. The graph-feature stack outperforms DPT on these transition-region tasks. Palantir entropy, CellRank fate uncertainty, and related transition scores were not evaluated and remain important comparisons; DPT does not represent all trajectory methods.

Future work should include: strictly inductive evaluation where test cells cannot influence each other or training features; donor-level or external-cohort validation; direct comparison with trajectory-method transition scores; independent marker-level biological validation; and a completed lung benchmark.

## Supporting information

Supplementary Material: Mathematical details, robustness checks, negative controls, sensitivity analyses, and support tables

## Code and Data Availability

All code, configurations, and result artifacts are available at https://github.com/ChimdiWalter/ricci_curvature. Datasets are loaded via standard API routes: Paul15 through Scanpy, pancreas and zebrafish through CellRank. The complete preprocessing, graph construction, curvature computation, and evaluation pipeline is documented in the repository README.

## 10 Conclusion

Discrete curvature provides dataset- and classifier-dependent incremental signal for some transition-region ranking tasks. The largest canonical Ollivier gain occurs on the pancreas Fev+ benchmark (five-split paired mean +0.092, 95% CI [0.076, 0.114]), but the primary evaluation is transductive, strict out-of-sample results are mixed, and matched marker analyses do not show independent biological enrichment beyond pseudotime and graph topology. Curvature-only models remain weak, and strong graph features are often sufficient.

