## Supplementary Material: Mathematical details, robustness checks, negative controls, sensitivity analyses, and support tables for "Dataset-Dependent Utility of Discrete Ricci Curvature for Transition-Region Ranking in Single-Cell Lineage Graphs"

#### 1 Supplement Overview

This supplement provides mathematical details, biological interpretation boundaries, robustness checks, additional analyses, support tables, and an artifact manifest.

| Supplement section | Role in review | Claim status |
|---|---|---|
| Mathematical Details | Defines graph construction, curvature, aggregation, residualization, and evaluation logic | Supports main methods claims |
| Biological Interpretation Details | Bounds the `Fev+`, zebrafish, bottleneck, and transfer interpretations | Supports main biological claim boundaries |
| Stricter Split Checks | Tests the zebrafish early-lineage conclusion under sample-token holdout | Appendix strengthening |
| Bottleneck Proxy Analyses | Reports the biologically motivated pancreas bottleneck proxy | Exploratory only |
| Transfer Analyses | Reports pairwise cross-dataset transfer asymmetry | Exploratory only |
| Negative Controls And Sensitivity | Shows random, rewired, and graph-construction checks | Supports robustness boundaries |
| Support Tables | Provides full metrics, paired deltas, supported-claim mapping, and validity boundaries | Main, appendix, exploratory, and future-work map |
| Lung No-Go Note | Explains why lung is excluded from current evidence | Future work only |

Table 1: Navigation guide for the supplement. Sections are organized by claim role so that main support, appendix strengthening, exploratory checks, and future-work boundaries remain separate.

#### 2 Mathematical Details

##### 2.1 Graph construction notation

For each dataset, cells are represented as nodes $V$ in a weighted undirected graph $G = (V, E, w)$. The canonical graph is rebuilt from PCA coordinates by Euclidean $k$-nearest-neighbor search with $k = 15$. If $d(u, v)$ is the PCA-space distance between neighboring cells, the heat-kernel weight is

$$w_{uv} = \exp\left(-\frac{d(u,v)^2}{2\sigma^2}\right),$$

where $\sigma$ is the median positive neighbor distance. The edge table stores both `weight` and `distance`; the latter is used by the transport computation for Ollivier-Ricci curvature.

#### 2.2 Forman-Ricci curvature

The implemented Forman-Ricci curvature is an edge statistic with unit node weights:

$$F(u,v) = w_{uv}\left[\frac{2}{w_{uv}} - \sum_{x\in N(u)\setminus\{v\}} \frac{1}{\sqrt{w_{uv}w_{ux}}} - \sum_{y\in N(v)\setminus\{u\}} \frac{1}{\sqrt{w_{uv}w_{vy}}}\right].$$

The statistic is treated as an empirical descriptor of local graph expansion and compression. No formal guarantee is claimed that this value identifies biological branch points.

#### 2.3 Ollivier-Ricci curvature

For each node $u$, the implementation defines a lazy neighborhood measure

$$\mu_u = \alpha\delta_u + (1-\alpha)\sum_{x\in N(u)} \frac{w_{ux}}{\sum_{z\in N(u)} w_{uz}}\delta_x,$$

with idleness $\alpha = 0.5$. For an edge $(u,v)$, Ollivier-Ricci curvature is

$$\kappa(u,v) = 1 - \frac{W_1(\mu_u,\mu_v)}{d_G(u,v)}.$$

Here $W_1$ is solved as an earth-mover distance between finite local measures, and $d_G$ is the graph distance induced by stored edge distances.

#### 2.4 Node-level aggregation

Curvature is computed on edges and then aggregated to cells. For each node and each curvature family, incident edge values are summarized by mean, minimum, maximum, and standard deviation. These summaries form the curvature-only feature tables and are concatenated with graph features for hybrid families.

#### 2.5 Hybrid feature construction

The evaluated families are graph features alone, Forman curvature only, Ollivier curvature only, graph plus Forman, graph plus Ollivier, random matched-dimension controls, and rewired graph controls. Curvature-only models answer whether curvature is sufficient by itself. The main scientific comparison is hybrid versus graph topology, because the relevant question is whether curvature adds information beyond an already strong graph representation.

#### 2.6 Residualization and anti-hub controls

Residualized curvature checks regress curvature summaries against graph-topology quantities such as degree and betweenness-style centrality before evaluation. These checks address a simple hubness alternative explanation. They are empirical controls, not proofs that curvature is a biological mechanism.

#### 2.7 Evaluation metrics

Branch-region tasks use exact-region AUPRC as the primary metric, with top-k and tolerant-region metrics as diagnostics. Fate-style tasks use macro-F1 to avoid hiding minority-class behavior. The pancreas and zebrafish fate-style tasks are intentionally narrow and label-supported.

##### 2.8 Bootstrap intervals and paired deltas

The reported confidence intervals are bootstrap intervals over repeated split-level summaries in the materialized artifacts. Paired deltas compare a left feature family against the graph-feature stack on the same task, graph setting, and split scaffold. This paired logic is why the manuscript emphasizes delta intervals against graph topology rather than only absolute scores.

##### 2.9 Empirical rather than theoretical status

The repository establishes empirical support, not a theorem. The supported conclusion is that curvature-derived node features can add controlled, dataset-dependent signal on some transition-region tasks. Exact biological branch-point localization, donor-level validation, and fully harmonized transfer remain future work.

#### 3 Biological Interpretation Details

##### 3.1 Pancreas Fev+ transition-region interpretation

The pancreas branch task is anchored on the coarse `Fev+` population. The task-design artifacts identify this state as a high-mixing endocrine transition region connected to beta, alpha, epsilon, delta, and upstream `Ngn3 high EP` neighborhoods. This makes `Fev+` the strongest annotation-backed branch-region proxy in the current package.

The main biological statement is that graph-plus-Ollivier features improve prioritization of this transition region over graph topology alone. The statement is not that the method discovers exact biological branch points.

##### 3.2 What branch-region ranking means biologically

Branch-region ranking is a transition-neighborhood task. A high score means that a model prioritizes cells in an annotation-informed region where lineage neighborhoods mix. The positive labels are proxies derived from annotations and graph-neighborhood structure. They are useful for controlled evaluation, but weaker than independent branch-point labels.

##### 3.3 Why this is not exact branch-point localization

Exact branch-point localization would require independently curated biological labels for individual branch cells or an external validation design. The current datasets provide transition-region annotations, lineage labels, stage windows, and graph structure. They do not provide exact branch-point ground truth.

##### 3.4 Zebrafish branch versus early-lineage interpretation

Zebrafish supports two different tasks. The branch-window ranking task asks whether models prioritize cells around the early stage transition window. The early-lineage task asks whether early cells can be classified as `Notochord` or `Prechordal Plate`. Curvature hybrids help on the former but not on the canonical version of the latter. That distinction is central to the paper.

##### 3.5 Why non-uniform behavior is plausible

Dataset behavior can differ because annotation granularity, developmental timing, graph density, lineage separability, and protocol structure differ. A local transition-region task can expose curvature signal, while a broad lineage task may already be solved by topology.

##### 3.6 Why curvature may help transition-region tasks

Curvature summarizes local edge-neighborhood structure. Transition regions often sit near local graph reorganization, where neighborhoods change composition and edge geometry may encode information not fully captured by scalar centrality alone. This is a biological plausibility argument, not a causal proof.

##### 3.7 Bottleneck proxy boundary

The pancreas late-`Ngn3 high EP` bottleneck proxy is biologically motivated and materialized, but the signal is weak and the label remains a proxy. It is therefore appendix-level exploratory evidence.

##### 3.8 Transfer asymmetry boundary

Pairwise transfer is asymmetric: some Paul15-pancreas directions transfer moderately, while transfers involving zebrafish remain weak. This matters because it prevents overclaiming and shows that feature portability depends on harmonized task semantics. Transfer remains exploratory.

#### 4 Stricter Split Checks

The zebrafish single-stage sample-token holdout is the strictest additional split check currently materialized. It restricts evaluation to the `05.3-50%` stage and holds out sample tokens rather than using only the canonical repeated node holdout. Graph topology remains strongest: the graph-feature stack reaches macro-F1 0.711, compared with 0.670 for graph-plus-Forman and 0.674 for graph-plus-Ollivier. This supports the main paper's conclusion that topology can already be sufficient for some early-lineage settings.

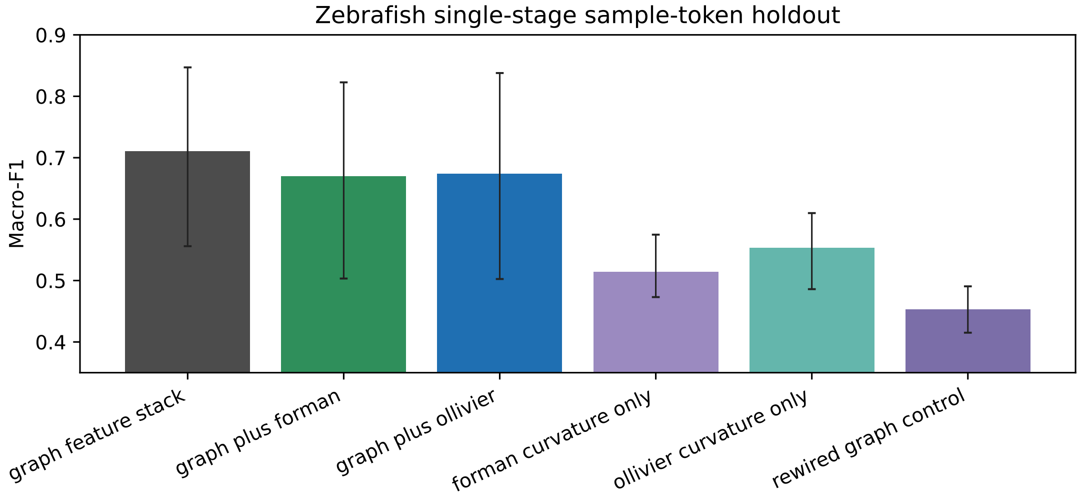


Figure 1: Zebrafish single-stage sample-token holdout. This is a stricter appendix check, not donor-level validation.

### 5 Bottleneck Proxy Analyses

The pancreas bottleneck proxy focuses on late-`Ngn3 high EP` cells and is biologically motivated, but it remains weak and proxy-based. Graph-plus-Ollivier reaches exact AUPRC 0.109 versus 0.096 for graph topology. This is not strong enough for a headline bottleneck detection claim.

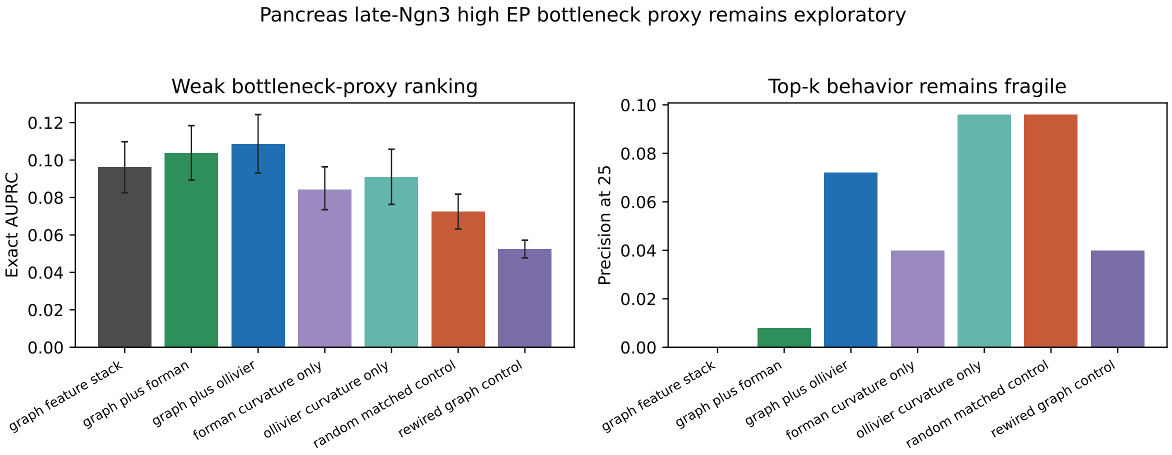


Figure 2: Pancreas bottleneck proxy results. The analysis is useful preparation for future biological bottleneck labels, but remains exploratory in the present evidence hierarchy.

### 6 Transfer Analyses

Pairwise transition-region transfer is materialized but asymmetric. The strongest directions are not sufficient to establish a transfer-first result, and transfers involving zebrafish remain weak. The transfer matrix is therefore interpreted as evidence of dataset dependence rather than broad portability.

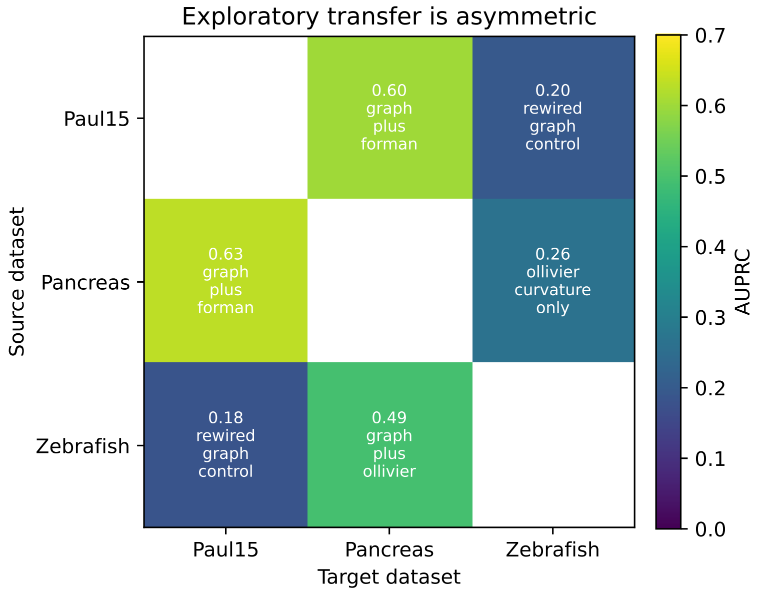


Figure 3: Exploratory pairwise transfer. Values show the best AUPRC among evaluated feature families for each source-target direction. Asymmetry keeps transfer appendix-level.

### 7 Negative Controls And Sensitivity

Random matched-dimension controls and rewired controls stay below the graph-feature stack on branch-region tasks. Graph-construction sensitivity affects exact magnitudes, but the branch-region pattern remains consistent with the main evidence structure.

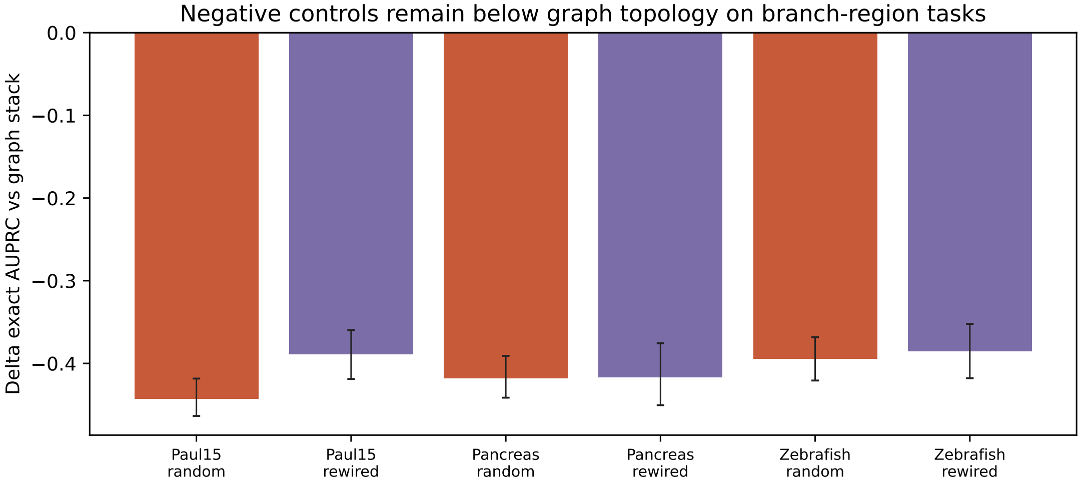


Figure 4: Negative controls on branch-region tasks. Controls remain below graph topology, supporting the view that the hybrid gains are not merely random feature expansion.

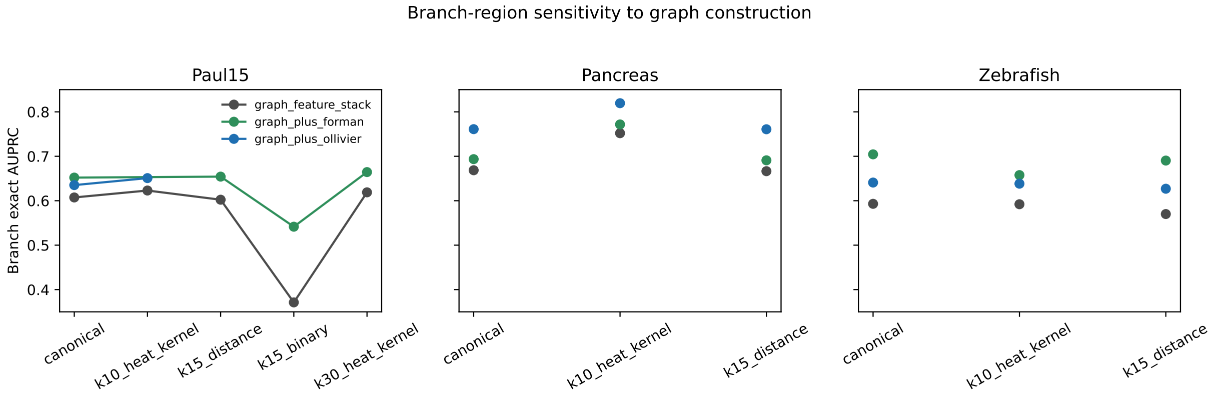


Figure 5: Branch-region sensitivity to graph construction. The exact magnitudes depend on graph settings, but the main conclusion remains hybrid and dataset-dependent.

#### 8 Support Tables

| Dataset | Task | Feature family | Metric | Value [CI] |
|---|---|---|---|---|
| Paul15 | branch_region_ranking | graph_feature_stack | auprc_exact | 0.607 [0.586, 0.628] |
| Paul15 | branch_region_ranking | forman_curvature_only | auprc_exact | 0.469 [0.448, 0.490] |
| Paul15 | branch_region_ranking | ollivier_curvature_only | auprc_exact | 0.272 [0.262, 0.283] |
| Paul15 | branch_region_ranking | graph_plus_forman | auprc_exact | 0.652 [0.627, 0.677] |
| Paul15 | branch_region_ranking | graph_plus_ollivier | auprc_exact | 0.635 [0.620, 0.650] |
| Paul15 | branch_region_ranking | random_matched_control | auprc_exact | 0.164 [0.158, 0.170] |
| Paul15 | branch_region_ranking | rewired_graph_control | auprc_exact | 0.219 [0.204, 0.228] |
| Pancreas | branch_region_ranking | graph_feature_stack | auprc_exact | 0.669 [0.637, 0.695] |
| Pancreas | branch_region_ranking | forman_curvature_only | auprc_exact | 0.305 [0.291, 0.319] |
| Pancreas | branch_region_ranking | ollivier_curvature_only | auprc_exact | 0.281 [0.264, 0.299] |
| Pancreas | branch_region_ranking | graph_plus_forman | auprc_exact | 0.694 [0.660, 0.724] |
| Pancreas | branch_region_ranking | graph_plus_ollivier | auprc_exact | 0.761 [0.729, 0.792] |
| Pancreas | branch_region_ranking | random_matched_control | auprc_exact | 0.250 [0.243, 0.258] |
| Pancreas | branch_region_ranking | rewired_graph_control | auprc_exact | 0.252 [0.237, 0.268] |
| Pancreas | preterminal_endocrine_fate_prediction | graph_feature_stack | macro_f1 | 0.594 [0.571, 0.627] |
| Pancreas | preterminal_endocrine_fate_prediction | forman_curvature_only | macro_f1 | 0.491 [0.470, 0.512] |
| Pancreas | preterminal_endocrine_fate_prediction | ollivier_curvature_only | macro_f1 | 0.456 [0.434, 0.479] |
| Pancreas | preterminal_endocrine_fate_prediction | graph_plus_forman | macro_f1 | 0.693 [0.671, 0.717] |
| Pancreas | preterminal_endocrine_fate_prediction | graph_plus_ollivier | macro_f1 | 0.676 [0.660, 0.699] |
| Zebrafish | branch_region_ranking | graph_feature_stack | auprc_exact | 0.593 [0.567, 0.620] |
| Zebrafish | branch_region_ranking | forman_curvature_only | auprc_exact | 0.337 [0.320, 0.356] |
| Zebrafish | branch_region_ranking | ollivier_curvature_only | auprc_exact | 0.334 [0.327, 0.345] |
| Zebrafish | branch_region_ranking | graph_plus_forman | auprc_exact | 0.704 [0.671, 0.744] |
| Zebrafish | branch_region_ranking | graph_plus_ollivier | auprc_exact | 0.641 [0.613, 0.669] |
| Zebrafish | branch_region_ranking | random_matched_control | auprc_exact | 0.199 [0.191, 0.206] |
| Zebrafish | branch_region_ranking | rewired_graph_control | auprc_exact | 0.208 [0.196, 0.220] |
| Zebrafish | early_lineage_prediction | graph_feature_stack | macro_f1 | 0.771 [0.755, 0.788] |
| Zebrafish | early_lineage_prediction | forman_curvature_only | macro_f1 | 0.564 [0.522, 0.595] |
| Zebrafish | early_lineage_prediction | ollivier_curvature_only | macro_f1 | 0.531 [0.495, 0.567] |
| Zebrafish | early_lineage_prediction | graph_plus_forman | macro_f1 | 0.764 [0.739, 0.790] |
| Zebrafish | early_lineage_prediction | graph_plus_ollivier | macro_f1 | 0.742 [0.714, 0.769] |

Table 2: Full canonical metric summary used by the submission. The machine-readable supplement asset bundle contains the same rows.

| Dataset | Task | Family | Metric | Delta vs graph [CI] |
|---|---|---|---|---|
| Paul15 | branch_region_ranking | graph_plus_ollivier | auprc_exact | 0.028 [0.021, 0.034] |
| Paul15 | branch_region_ranking | graph_plus_forman | auprc_exact | 0.045 [0.037, 0.051] |
| Paul15 | branch_region_ranking | random_matched_control | auprc_exact | -0.443 [-0.464, -0.418] |
| Paul15 | branch_region_ranking | rewired_graph_control | auprc_exact | -0.389 [-0.419, -0.360] |
| Paul15 | branch_region_ranking | graph_plus_ollivier | auroc_exact | 0.007 [0.005, 0.009] |
| Paul15 | branch_region_ranking | graph_plus_forman | auroc_exact | 0.007 [0.005, 0.009] |
| Paul15 | branch_region_ranking | random_matched_control | auroc_exact | -0.426 [-0.443, -0.411] |
| Paul15 | branch_region_ranking | rewired_graph_control | auroc_exact | -0.343 [-0.371, -0.314] |
| Paul15 | branch_region_ranking | graph_plus_ollivier | precision_at_k_25_exact | 0.016 [0.000, 0.032] |
| Paul15 | branch_region_ranking | graph_plus_forman | precision_at_k_25_exact | 0.048 [-0.008, 0.096] |
| Paul15 | branch_region_ranking | random_matched_control | precision_at_k_25_exact | -0.608 [-0.696, -0.528] |
| Paul15 | branch_region_ranking | rewired_graph_control | precision_at_k_25_exact | -0.424 [-0.504, -0.360] |
| Paul15 | branch_region_ranking | graph_plus_ollivier | auprc_tolerant | 0.002 [0.001, 0.004] |
| Paul15 | branch_region_ranking | graph_plus_forman | auprc_tolerant | 0.003 [-0.000, 0.005] |
| Paul15 | branch_region_ranking | random_matched_control | auprc_tolerant | -0.533 [-0.547, -0.514] |
| Paul15 | branch_region_ranking | rewired_graph_control | auprc_tolerant | -0.498 [-0.525, -0.475] |
| Pancreas | branch_region_ranking | graph_plus_forman | auprc_exact | 0.025 [0.012, 0.036] |
| Pancreas | branch_region_ranking | graph_plus_ollivier | auprc_exact | 0.092 [0.076, 0.114] |
| Pancreas | branch_region_ranking | random_matched_control | auprc_exact | -0.418 [-0.441, -0.391] |
| Pancreas | branch_region_ranking | rewired_graph_control | auprc_exact | -0.417 [-0.451, -0.376] |
| Pancreas | preterminal_endocrine_fate_prediction | graph_plus_forman | macro_f1 | 0.099 [0.075, 0.125] |
| Pancreas | preterminal_endocrine_fate_prediction | graph_plus_ollivier | macro_f1 | 0.082 [0.070, 0.093] |
| Pancreas | preterminal_endocrine_fate_prediction | random_matched_control | macro_f1 | [, ] |
| Pancreas | preterminal_endocrine_fate_prediction | rewired_graph_control | macro_f1 | [, ] |
| Zebrafish | branch_region_ranking | graph_plus_forman | auprc_exact | 0.111 [0.090, 0.128] |
| Zebrafish | branch_region_ranking | graph_plus_ollivier | auprc_exact | 0.048 [0.041, 0.055] |
| Zebrafish | branch_region_ranking | random_matched_control | auprc_exact | -0.394 [-0.421, -0.368] |
| Zebrafish | branch_region_ranking | rewired_graph_control | auprc_exact | -0.385 [-0.418, -0.352] |
| Zebrafish | early_lineage_prediction | graph_plus_forman | macro_f1 | -0.007 [-0.017, 0.005] |
| Zebrafish | early_lineage_prediction | graph_plus_ollivier | macro_f1 | -0.030 [-0.058, 0.001] |
| Zebrafish | early_lineage_prediction | random_matched_control | macro_f1 | [, ] |
| Zebrafish | early_lineage_prediction | rewired_graph_control | macro_f1 | [, ] |

Table 3: Paired deltas against the graph-feature stack on the canonical graph.

| Level | Claim | Artifact |
|---|---|---|
| main supported | Curvature adds useful hybrid signal on some branch-region tasks. | cross-dataset summary table |
| main supported | Pancreas Fev+ branch ranking is the clearest Ollivier-positive result. | pancreas Phase 3 summary and paired-delta tables |
| main supported | Zebrafish early-lineage prediction is graph-topology dominated on the canonical graph. | zebrafish Phase 3 summary and paired-delta tables |
| appendix strengthening | Zebrafish sample-token holdout still favors graph topology. | zebrafish sample-token holdout summary |
| exploratory | Pancreas late-Ngn3 high EP bottleneck proxy is biologically grounded but weak. | pancreas bottleneck proxy summary |
| exploratory | Pairwise transfer is asymmetric and not a headline success claim. | pairwise transfer summary |
| future work only | Lung is downloaded and audited but not included as completed evidence. | lung no-go decision record |

Table 4: Supported-claims mapping. Main, appendix, exploratory, and future-work claims are tied to concrete artifacts.

| Boundary | Scope | Manuscript policy |
|---|---|---|
| Branch-region labels are proxies | Paul15, pancreas, zebrafish branch tasks | say transition-region ranking, not exact branch-point localization |
| Pancreas fate is preterminal | Pancreas fate task | do not call it earliest endocrine commitment |
| Zebrafish early lineage is canonical non-gain | Zebrafish early-lineage task | present as graph-topology sufficiency, not as failed experiment |
| Bottleneck remains exploratory | Phase 4 pancreas bottleneck proxy | appendix only; no biological bottleneck detection claim |
| Transfer remains exploratory | Phase 4 pairwise transfer | appendix only; no transfer-first narrative |
| No donor-level validation | all supported tasks | state as limitation and future direction |

Table 5: Limitations and validity boundaries that govern manuscript language.

| Analysis | Headline number | Interpretation | Claim level |
|---|---|---|---|
| zebrafish sample-token holdout | graph macro-F1 0.711; graph+Forman 0.670 | stricter split still favors graph topology | appendix strengthening |
| pancreas bottleneck proxy | graph AUPRC 0.096; graph+Ollivier 0.109 | biologically grounded but weak proxy | exploratory |
| pairwise transfer | best AUPRC 0.630 (pancreas to paul15) | asymmetric; not a headline transfer claim | exploratory |
| pancreas palantir reference | branch AUPRC 0.184 | narrow reference baseline is weaker than graph stack | appendix strengthening |

Table 6: Appendix extension summary. These analyses sharpen the main story but do not become headline claims.

### 9 Lung No-Go Note

The official CellRank lung dataset is downloaded and schema-audited, but no stable benchmark artifact is present in the current evidence package. Lung is therefore future work only in this submission. A future upgrade would require a documented lung task design, materialized label files, model-family metrics, sensitivity checks, and biological interpretation notes.

### 10 Supplement Manifest

The machine-readable supplement manifest lists generated tables, figures, copied appendix assets, and source evidence policy. It is produced by the build scripts and is described in the reproducibility notes.
